# Hepatocyte Estrogen Receptor α Mediates Estrogen-induced Augmentation of Hepatic Mitochondrial Respiration Following Ovariectomy

**DOI:** 10.64898/2026.02.22.706993

**Authors:** Edziu Franczak, Benjamin A. Kugler, Sebastian F. Salathe, Julie A. Allen, Colin S. McCoin, E. Matthew Morris, John P. Thyfault

## Abstract

Whole-body estrogen receptor α (ERα) knockout mice develop hepatic steatosis; however, liver-specific ERα knockout (LERKO) mice fail to recapitulate this susceptibility and maintain normal hepatic mitochondrial function. However, estrogen-mediated protection against hepatic steatosis is lost in LERKO mice following ovariectomy (OVX). Here, we tested whether loss of hepatic ERα blunts estrogen modulation of hepatic mitochondrial respiratory capacity and mitochondrial proteome following ovariectomy (OVX). Sham or ovariectomy (OVX) surgery was performed in middle-aged female mice (36-40 weeks), followed by AAV injection to generate Control (Con; GFP) or LERKO mice (Cre). All mice were placed on a high-fat diet (HFD) for 10 weeks following surgery. Half of the OVX mice received 17-beta estradiol (E2) replacement (OVX+E2) for the last 4 weeks of HFD. OVX mice had greater body mass and adiposity, which was reversed by E2 replacement in both Con and LERKO mice. While E2 replacement reduced steatosis in both Con and LERKO OVX mice, the LERKO OVX mice maintained greater hepatic triglyceride content. E2 replacement promoted greater basal and ADP-stimulated (State 3) mitochondrial respiration in Con OVX but not in LERKO OVX mice under palmitate-supported conditions. Changes in mitochondrial respiration could not be attributed to altered responses to changes in energy demand (G_ATP_) or to alterations in mitochondrial H_2_O_2_ production.

Conversely, maximal coupled branched-chain amino acid-supported respiration was universally suppressed by E2 replacement. Proteomics analysis revealed E2-mediated reductions in hepatic mitochondrial energy transduction, with relatively minimal differences between Con and LERKO mice. In conclusion, post-ovariectomy estrogen treatment reduces steatosis in the absence of hepatic ERα; however, triglyceride levels remain higher, and mitochondrial respiratory deficits persist despite similar proteomic signatures, suggesting that ERα signaling is required for optimal estrogen hepatic responsiveness.

## Introduction

In women, the primary site of estrogen synthesis occurs within the ovaries through aromatase activity^1^. 17β-estradiol (E2) serves as the most prominent form of estrogen in premenopausal women, producing cellular responses through the activation of various estrogen receptors, including estrogen receptor (ER) α, ERβ^2,3^, and G-protein coupled receptor 1 (GPER1)^4^. In the classical E2 signaling pathway, nuclear ERs (ERα, ERβ) exert transcriptional control within the nucleus by binding to estrogen-responsive elements (EREs) located throughout the genome or by modulating the activation of various other transcription factors^5^. The liver is considered one of the most sexually dimorphic tissues, with >70% of hepatic genes/proteins displaying sexually divergent expression in both humans and rodents^6,7^. Within the liver, ERα serves as the primary estrogen receptor, capable of binding to approximately 5,568 unique sites within the hepatic genome^8^, exerting extensive transcriptional control over genes involved in fatty acid oxidation, cholesterol/bile acid metabolism, amino acid catabolism, and mitochondrial electron transport system (ETS)^8–13^. Recent use of liver-specific ERα knockout mice (LERKO) has revealed ∼32% of all sex-biased gene expression profiles within the liver are directly regulated by ERα^11^.

Despite the transcriptional regulation imposed by ERα, the role of E2 signaling in promoting innate protection against metabolic dysfunction-associated steatotic liver disease (MASLD) in women remains unknown^14^.

Women display lower rates of MASLD compared to men, but following the onset of menopause and the subsequent depletion of endogenous E2 production, innate protection against the development of hepatic steatosis is completely ablated in women^15–17^. In rodents, surgical (ovariectomy; OVX)^18^ and chemical-induced (Vinyl cyclohexene dioxide; VCD) menopause^19^ promotes increased propensity for intrahepatic lipid accumulation^18^. In both women and rodent models, estrogen hormone replacement therapy (HRT) protects against or reduces hepatic steatosis following loss of ovarian function^18,20–22^. Thus far investigations cast doubt on the central role of hepatic ERα in modulating this protection against hepatic steatosis because of conflicting results in ovarian-intact mice^9,13,23,24^. However, recent reports have shown that following OVX, E2 replacement requires expression of hepatic ERα to effectively reduce liver triglycerides^21,22^, suggesting a unique role of ERα in modulating hepatic lipid handling after the loss of ovarian function.

In human subjects, MASLD is associated with dynamic alterations in hepatic mitochondrial respiratory capacity, characterized by initial compensatory increases in mitochondrial respiration during the development of hepatic steatosis, followed by a decline in respiratory capacity with liver injury and significant disease progression^25^. We have recently reported intrinsic sex differences in hepatic mitochondrial function, with ovarian-intact female mice possessing heightened mitochondrial respiratory capacity and reduced mitochondrial H_2_O_2_ production compared to males^26^. Importantly, metabolic stressors (i.e. high fat diet and exercise) further intensify these differences in mitochondrial respiration, with ovarian-intact female mice uniquely capable of expanding mitochondrial respiratory capacity, while consistently retaining lower mitochondrial H_2_O_2_ emission^27,28^. While these mitochondrial phenotypes may serve to protect the female liver from the development of hepatic steatosis^18^, limited investigation exists regarding the direct role of E2 in modulating hepatic mitochondrial function. We previously found that mice with liver specific ERα knockouts (LERKO) displayed no mitochondrial phenotypes or increases in steatosis in ovary intact females, despite some evidence of differences in hepatic transcriptional regulation^29^. Herein, we tested the hypothesis that E2-mediated resolution of high-fat diet-induced hepatic steatosis following OVX requires ERα-dependent stimulation of hepatic mitochondrial respiration. Our findings suggest that E2 is capable of effectively reducing hepatic steatosis and mitochondrial H_2_O_2_ production independent of ERα signaling. However, subsequent alterations in mitochondrial respiratory capacity necessitate ERα expression within hepatocytes despite relatively limited proteomic alterations directly modulated by ERα. These findings implicate ERα as a critical regulator of mitochondrial function within the liver following the loss of ovarian function suggesting it is a critical therapeutic target for pharmaceutical treatment of MASLD in post-menopausal women.

## Methods

### Experimental Treatment

Female homozygous ERα floxed mice (ERα^fl/fl^) that were 36-40 weeks of age were utilized in the present study, with genotype confirmed via polymerase chain reaction (PCR)-based genotyping of tail snips. Mice underwent either a sham or bilateral OVX surgery to deplete systemic sex hormone production. Immediately following surgical procedure, OVX mice were injected with either an adeno-associated virus containing either Cre (AAV.TBG.PI.Cre.rBG) to generate hepatocyte-specific deletion of ERα expression (LERKO), or green fluorescent protein (GFP) (pAAV.TBG.PI.eGFP.WPRE.bGH) to retain expression of floxed regions (Con) (n = 22 per treatment group), as previously described^9^. Sham mice received an injection of AAV containing GFP. Mice were then transitioned to high fat diet (HFD; Research diets D12451; 45%kcal fat, 17% kcals sucrose, 1% cholesterol wt./wt.; 4.68 kcal/g) for 10 weeks. All mice were co-housed at thermoneutral temperature (30°C) on a reverse light cycle (dark: 10:00-22:00) with *ad libitum* access to water and HFD.

### Estrogen Replacement

Following the initial 6 weeks of dietary intervention, OVX mice underwent surgical implantation of 20mM silastic tubing (3 mM surgical glue capping) between the scapulae blades containing sesame oil or E2 replacement (36μg 17β-estradiol/mL sesame oil), as previously described^30^. Silastic tubing was surgically replaced after 2 weeks to prevent E2 depletion. All sham mice received SO treatment.

### Anthropometrics and Tissue collection

Body weight was measured weekly throughout the study. Immediately prior to initial E2 replacement surgery and at the end of the dietary intervention, body composition was assessed via magnetic resonance imaging (EchoMRI, Houston, TX). Fat free mass was determined by subtracting fat mass from total body mass.

Vaginal cytology was performed to determine estrous stage immediately prior to euthanasia, as previously described^31^. All mice were anesthetized using phenobarbital (0.5mg/g BW) before terminal procedure following a 2 hour fast (7:00am – 9:00 am). Blood was immediately collected via cardiac puncture and left a room temperature to clot before being placed on ice for 10 minutes. Blood was then centrifuge at 7,000 x g (10 min, 4°C) and serum was separated. Livers were quickly excised and flash frozen in liquid nitrogen for later processing, while other portions were saved for histology and mitochondrial isolation. Weight wets for Adipose tissue (perigonadal and inguinal) and uterus were recorded.

### Liver Triglycerides

500 μg protein from frozen liver tissue was used for lipid extraction via the Folch method, before lipids were resuspended in 100μL of tert-butanol-Triton X solution, as previously described^32^.

Liver triglyceride content was measured using the commercially available kit (TR0100, Sigma Aldrich).

### Western blot Protein Quantification

80-100 mg of pulverized liver tissue were used to quantify protein expression as previously described^9^. SDS-PAGE was used to separate proteins before transferring to polyvinylidene difluoride (PVDF) membrane. Membranes were probed ERα to confirm hepatocyte deletion (Abcam ab32063). Membranes were imaged, and densitometry was used to quantify protein bands (Bio-Rad Laboratories, Hercules, CA). All protein bands were normalized to total protein using 0.1% amido-black staining.

### Hepatic Mitochondrial Isolation

Following excision, a portion of the liver was placed in 8 mL of ice-cold mitochondrial isolation buffer (220 mM mannitol, 70 mM sucrose, 10 mM Tris, 1 mM EDTA, pH at 7.4) and homogenized using a Teflon pestle. Dounce glass-on-glass homogenization in combination with density centrifugation was performed as previously described to generate a mitochondrial enriched fraction^32^. Briefly, liver homogenate was centrifuged at 1,500 x g (10 min at 4°C) and the supernatant filtered through cheese cloth before being pelleted (8,000 x g, 10min at 4°C).

The pellet was then resuspended in 6 mL of mitochondrial isolation buffer, before being pelleted again at 6,000 x g (10min at 4°C). Following resuspension in 4 mL of mitochondrial isolation buffer containing 0.1% fatty acid-free BSA, mitochondrial fractions were spun at 4,000 x g (10min at 4°C). The final isolated mitochondrial pellet was resuspended in 350-400 μL of modified MiR05 mitochondrial respiration buffer (0.5 mM EGTA, 3 mM MgCl_2_, 60 mM KMES, 20 mM glucose, 10 mM KH_2_PO_4_, 20 mM HEPES, 110 mM sucrose, and 0.1% fatty acid-free BSA, pH adjusted to 7.1). Bicinchoninic acid assay was performed to determine protein concentration of the isolated mitochondrial fraction.

### Mitochondrial Respiration

Real-time mitochondrial oxygen consumption (*J*O_2_; pmol s^-1^ mL ^-1^) and H_2_O_2_ flux (pmol s^-1^ mL^-1^) were measured simultaneously using the Oroboros O2k fluorometer (Oroboros Instruments, Innsbruck, Austria) as previously described^33^. Following air calibration, 100-150 μg of isolated hepatic mitochondria was added to the Oroboros chambers with appropriate substrate conditions. All mitochondrial respiration experiments were performed at 37°C in 2 mL of modified MiR05 respiration buffer with the addition of 2 mM malate, 63.5 μM free CoA, and 2.5 mM _L_-carnitine, continuously stirring at 750 rpm.

Palmitate-supported respiration was performed in the presence of 10 μM palmitoyl-CoA (PCoA) under basal/leak respiratory conditions. Coupled mitochondrial respiration was achieved following the addition of adenosine 5’-disphosphate (ADP; 2.5 mM). Palmitoyl-carnitine (PC; 10μM) and succinate (state 3S; 10 mM) were then added to determine Cpt1α-dependent and maximal complex I and II respiration, respectively. Maximal uncoupled respiration was achieved following serial titration of carbonyl cyanide-*p*-trifluoromethoxy phenylhydrazone (FCCP).

Percent (%) electron leak was determined by correcting H_2_O_2_ flux by *J*O_2_ under basal and state 3 respiratory conditions. Branched-chain keto-acid (BCKA) respiration was determined following the addition of 1 mM α-ketoisocaproic acid (KIC) and 1 mM potassium bicarbonate under basal conditions. State 3 respiration was determined following the addition of ADP (2.5 mM). α-ketoisovaleric acid (KIV) followed by α-keto-β-methylvaleric acid (KMV) were added to determine both KIV and KMV-dependent respiration. Coupling control ratios (CCR) were calculated for both PCoA- and BCKA-supported respiration by dividing basal *J*O_2_ rates by State 3 *J*O_2_ rates under both substrate conditions, as previously described^27,28,34^.

### Mitochondrial Antioxidant Capacity

Following the addition of PCoA (10 mM) and PC (10 mM) to the modified MiR05 respiration buffer, mitochondrial isolates were injected into the chambers of the Oroboros O2k system. H_2_O_2_ flux was measured under basal respiratory conditions (absence of ADP) to maximize H_2_O_2_ production rates. Peroxiredoxin antioxidant capacity was determined following the injection of Auranofin (AF; 0.25 μM), an inhibitor of the peroxiredoxin antioxidant pathway. Similarly, 1-chloro-2,4-dinitrobenzene (50μM CDNB) was then added to deplete reduced glutathione (GSH), to determine the antioxidant capacity within the glutathione antioxidant pathway.

### Creatine Kinase (CK) Clamp

A modified version of the creatine kinase clamp was performed to determine change in mitochondrial respiration under a physiological range of free energy of ATP hydrolysis (ΔG_ATP_), as previously described^35^. Following the addition of creatine monohydrate (5 mM), PCoA (10 μM), PC (10 μM), creatine kinase (CK; 20U), and adenosine 5’-triphosphate (2.5 mM) to the modified MiR05 respiration buffer, mitochondrial *J*O_2_ was measured following the addition of phosphocreatine (PCr) at concentration of 1 mM, 3 mM, 6 mM, 9 mM, 12 mM, 15 mM, 18 mM, 21 mM within the chamber. Mitochondrial conductance was determined by calculating the linear relationship between *J*O_2_ and ΔG_ATP_, as previously described^36^.

### Hepatic Proteomics via Orbitrap Astral DIA

Total protein from each sample was reduced, alkylated, and purified by chloroform/methanol extraction prior to digestion with sequencing grade modified porcine trypsin (Promega).

Tryptic peptides were then separated by a reverse phase Ion-Opticks-TS analytical column (25 cm x 75 um with 1.7 um C18 resin) supported by an EASY-Spray nano-source and stabilized with a Heater THOR Controller (Ion-Opticks) at 60°C. Peptides were trapped and eluted from a (PepMap Neo, 300um x 5mm Trap) using a Vanquish Neo UHPLC nano system (Thermo Scientific) which kept the samples at 11°C before injection. Peptides were eluted at a flow rate of 0.350uL/min using a 35 min gradient from 98% Buffer A:2% Buffer B to 94.5:5.5 at 0.1 minutes to 56:44 at 27.1 minutes followed by a column wash of 45:55 at 29.7 minutes to 1:99 at 35 minutes followed by equilibration back to 98:2. Eluted peptides were ionized by electrospray (2.5 kV) followed by mass spectrometric analysis on an Orbitrap Astral mass spectrometer (Thermo). Precursor spectra were acquired from 380-980 Th, 240,000 resolution, normalized AGC target 200%, maximum injection time 3 ms. DIA acquisition on the Orbitrap Astral was configured to acquire 199, 3 Th window from 380-980 Th, 25% HCD Collision Energy, normalized AGC target 100%, maximum injection time 3 ms. Fragment (MS2) scan range from 150-2000 Th with an RF Lens (%) set to 40.

## Data Analysis: Spectronaut – VSN and IPA

Following data acquisition, data were searched using Spectronaut (Biognosys version 19.5) against the UniProt *Mus musculus* database (Proteome ID: UP000000589, Taxon ID: 10090, 1^st^ version of 2025) using the directDIA method with an identification precursor and protein q-value cutoff of 1%, generate decoys set to true, the protein inference workflow set to maxLFQ, inference algorithm set to IDPicker, quantity level set to MS2, cross-run normalization set to false, and the protein grouping quantification set to median peptide and precursor quantity. Fixed Modifications were set to Carbamidomethyl (C) and variable modifications were set to Acetyl (Protein N-term), Oxidation (M). Protein MS2 intensity values were assessed for quality using ProteiNorm (Graw *et al.*). The data was normalized using VSN (Huber et al) and analyzed using proteoDA to perform statistical analysis using Linear Models for Microarray Data (limma) with empirical Bayes (eBayes) smoothing to the standard errors (Thurman et al, Ritchie et al). Proteins with an FDR adjusted p-value < 0.05 and a fold change > 2 were considered significant.

Ingenuity Pathway Analysis software (IPA, Qiagen) was used to perform pathway analysis for group comparisons. For assessment of the mitochondrial proteome, all identified proteins were cross-referenced against the MitoCarta3.0 database^37^. For assessment of individual protein expression across groups, the MS2 spectral abundance/intensity was utilized, followed by normalization to the summed MS2 abundance of all identified mitochondrial proteins, as previously described^34^.

### Statistical Analysis

Statistics were performed in SPSS Statistics 29 (IBM, Armonk, NY). Outliers were identified and removed using the Grubbs method using Prism 10 (GraphPad Software, San Diego, CA). A unpaired t-test was used to determine significant differences between Sham mice versus OVX or OVX+E2 mice, respectively. To determine differences between Con and LERKO OVX mice, a 2-Way ANOVA was performed (Con vs LERKO, OVX vs E2 Replacement). Post hoc analysis was conducted following the detection of a significant interaction between LERKO and E2 replacement using Fisher’s least significant difference (LSD) to assess significant pairwise comparisons. A post-hoc analysis was performed following detection of a main effect for proteomics datasets due to the limited samples size (n=5). Statistical significance was set at p < 0.05. For pathway analysis performed using IPA, significance was determined as a Z-score > ± 2 and a -log fold change of >1.3. All data are represented as means ± standard error (SEM) (GraphPad Prism 10).

## Results

### E2 replacement reduces adiposity and protects against hepatic steatosis

Figure 1A depicts the overall study design, including experimental groups and duration of OVX/sham and diet conditions. Following the initial 6 weeks of dietary intervention, Con OVX mice collectively possessed greater body mass (p<0.05), and fat mass (p < 0.05) compared to Sham mice (comparision not shown); however, this did not reach significance when Con OVX and Con OVX+E2 were assessed as separate groups (**Table 1**). No differences in body composition were observed between Con OVX and LERKO OVX mice. In response to E2 replacement, both Con and LERKO mice displayed sustained reductions in body mass (Fig. 1B, main effect p<0.001). Reductions in fat mass served as the primary contributor of decreases in body mass (**Table 1**, main effect p<0.001), with reductions in the mass of both perigonadal and inguinal fat pads (**Table 1**, main effect p<0.001). We identified no discernible differences in fat-free mass between groups. Body mass and composition were not different between sham and both Con OVX groups (+/- E2 replacement), although this is likely attributed to the age of the mice used in this study. Importantly, LERKO and Con mice did not differ in any measure of body composition. At the time of experimentation, Sham mice were effectively cycling through all four stages of the estrous cycle (Fig. 1C). However, all OVX mice (both control and LERKO) were stuck in diestrus, while all mice receiving E2 replacement were in proestrus (Fig. 1D**-E**).

**Figure 1.**
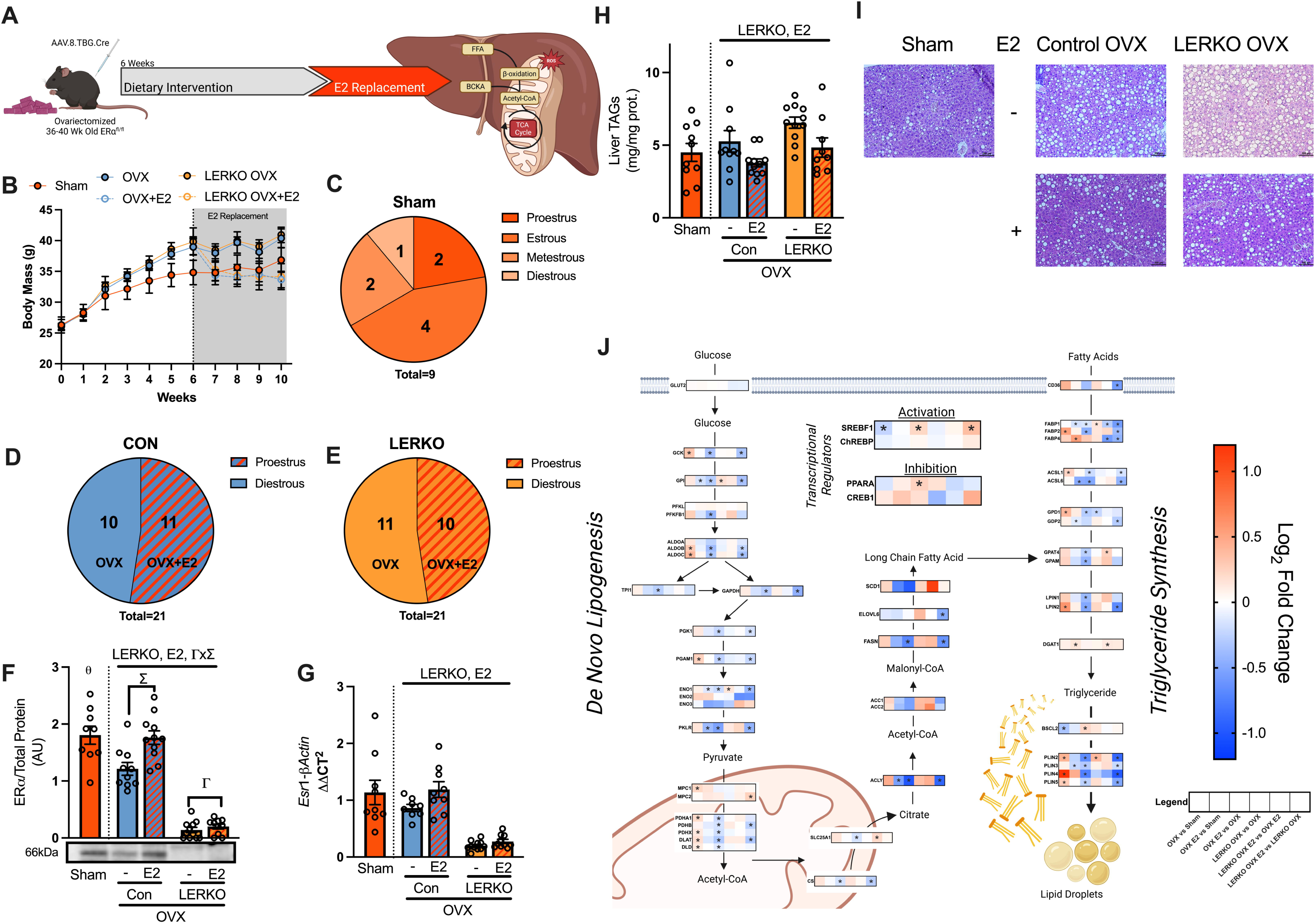
***E2 replacement restores body composition following OVX*.** Depiction of study timeline (A). Weekly change in body mass during 10-week intervention (B). Estrous stage for Sham (C), Con (D) and LERKO mice (E) were determined at time of experimentation. Hepatic ERα protein (F) and mRNA expression (G) were quantified to confirm effective knock-out of hepatocyte ERα. Liver triglycerides (H) and H and E staining (I) were performed to assess hepatic steatosis levels. Proteomic Log_2_ fold change was used to assess alterations in the expression of proteins involved in *de novo* lipogenesis and triglyceride synthesis (n=5 per group). Data in (B-H) are represented as means ± SE (n=9-11 per group). E2 and LERKO represents main effect of E2 replacement or hepatocyte ERα expression following Two-way ANOVA. Interaction between E2 replacement and LERKO are represented by εxΓ, ε representing significant post-hoc for E2-replacements, and Γ representing a significant post-hoc for LERKO. Data in (J) are log_2_ fold change comparison between individual groups. Significance values are p < 0.05.

**Table 1.** Anthropometrics

|  | Sham | Con |  | LERKO |  | 2-way Anova p-value |  |  |
| --- | --- | --- | --- | --- | --- | --- | --- | --- |
|  |  | OVX | OVX+E2 | OVX | OVX+E2 | Genotype | E2 Replacment | Interaction |
| Pre-E2 Replacement |  |  |  |  |  |  |  |  |
| Body Mass (g) | 34.82 ± 1.99 | 38.98 ± 1.32 | 39.07 ± 1.37 | 39.83 ± 0.79 | 39.99 ± 2.18 | 0.57 | 0.95 | 0.98 |
| Fat Mass (g) | 13.47 ± 1.65 | 17.47 ± 1.09 | 17.71 ± 1.11 | 18.19 ± 0.76 | 17.98 ± 1.65 | 0.78 | 0.88 | 0.91 |
| Fat-Free Mass (g) | 21.35 ± 0.55 | 21.51 ± 0.37 | 21.37 ± 0.44 | 21.64 ± 0.29 | 22.01 ± 0.61 | 0.19 | 0.99 | 0.83 |
| Post-E2 Replacement |  |  |  |  |  |  |  |  |
| Body Mass (g) | 36.86 ± 1.97 | 40.39 ± 1.52 | 33.68 ± 1.34 | 40.94 ± 1.25 | 34.15 ± 2.11 | 0.75 | <0.001 | 0.98 |
| Fat Mass (g) | 14.87 ± 1.68 | 18.52 ± 1.24 | 10.97 ± 1.17 | 18.78 ± 1.01 | 11.29 ± 1.65 | 0.79 | 0.001 | 0.16 |
| Fat-Free Mass (g) | 21.99 ± 0.49 | 21.87 ± 0.43 | 22.71 ± 0.38 | 22.15 ± 0.42 | 22.86 ± 0.54 | 0.63 | 0.09 | 0.89 |
| Perigonadal Fat (g) | 2.09 ± 0.25 | 2.59 ± 0.15 | 1.59 ± 0.17 | 2.73 ± 0.10 | 1.63 ± 0.21 | 0.56 | <0.001 | 0.74 |
| Inguinal Fat (g) | 1.25 ± 0.14 θ | 1.76 ± 0.14 | 0.83 ± 0.17 | 1.61 ± 0.12 | 0.98 ± 0.14 | 0.99 | <0.001 | 0.29 |
| Intrahepatic Triglycerides (g) | 4.50 ± 0.62 | 5.27 ± 0.74 | 3.78 ± 0.28 | 6.55 ± 0.39 | 4.55 ± 0.66 | <0.05 | <0.01 | 0.8351 |
| Uterine Mass (g) | 0.13 ± 0.02 θ,ε | 0.02 ± 0.003 | 0.19 ± 0.009 | 0.02 ± 0.002 | 0.21 ± 0.01 | 0.13 | <0.0001 | 0.18 |

Data are presented as means ± SE. n = 9-11/group. Bold type signifies significant p-value (< 0.05). θ and ε represents significant unpaired t-test between Sham and Con OVX and Sham Con OVX+E2, respectively. Con, Control; LERKO, hepatocyte-specific estrogen receptor α knockout; OVX, ovariectomy; E2, estradiol.

Similarly, uterine mass was used to further confirm OVX status and efficacy of E2 replacement (Fig. 1E, main effect OVX and E2 replacement p<0.001). ERα expression was assessed at both the proteomic (Fig. 1F) and transcriptional levels (Fig. 1G), confirming the effective knockout of hepatocyte ERα expression, with E2 replacement promoting increased ERα expression in both Con and LERKO OVX mice (p<0.0001). Residual ERα expression is likely attributed to retained expression in non-hepatocyte cells within the liver.

Recent reports have suggested that hepatic ERα directly modulates protection against hepatic steatosis post-OVX^21,22^. We next assessed measures of hepatic triglyceride content, revealing an expanded intrahepatic triglyceride content in LERKO versus Con mice (Fig. 1H**-I**), with a main effect (p < 0.05). However, E2 replacement promoted substantial reductions in hepatic triglyceride levels in both Con and LERKO OVX mice, suggesting hepatic ERα expression is not necessary for E2-mediated reductions in liver triglyceride levels (Fig. 1H**-I**, main effect p<0.01). Proteomics revealed that E2 treatment reduced proteins mediating *de novo* lipogenesis (DNL) and triglyceride synthesis, pathways regularly upregulated in hepatic steatosis, in both Con and LERKO groups (Fig. 1J). This would suggest that ERα-mediated signaling mechanisms play a relatively minimal role in modulating E2-stimulated reductions in lipid synthesis pathways underlying excessive hepatic triglyceride content.

### E2-mediated Alterations in the Hepatic Proteome

To decipher the distinct proteins under direct expressional control by E2, and more specifically, ERα-mediated regulation, we next performed untargeted proteomics analysis on whole liver tissue. Comparison analysis across our various experimental groups revealed several central pathways heavily influenced by E2-replacement, including upregulating mRNA transcription and processing and the immune response, while specifically reducing mitochondrial protein synthesis/degradation and processes involved in mitochondrial energy transduction (Fig. 2A). To determine whether any of these responses were directly attributed to ERα-mediated signaling events, we directly compared pathway analysis between LERKO OVX+E2 versus Con OVX+E2. We identified relatively few divergent pathways between these groups, with the largest differences being identified in mRNA processing, DNA base repair, mTOR signaling, and oxidative phosphorylation (OxPhos) (Fig. 2B, Log p>1.3, Z score > ±2). Intriguingly, OxPhos and its components (respiratory electron transport, Complex I biogenesis, and Complex IV assembly) were upregulated in LERKO OVX+E2 mice to a greater degree than the control OVX+E2mice (Fig. 2B).

**Figure 2.**
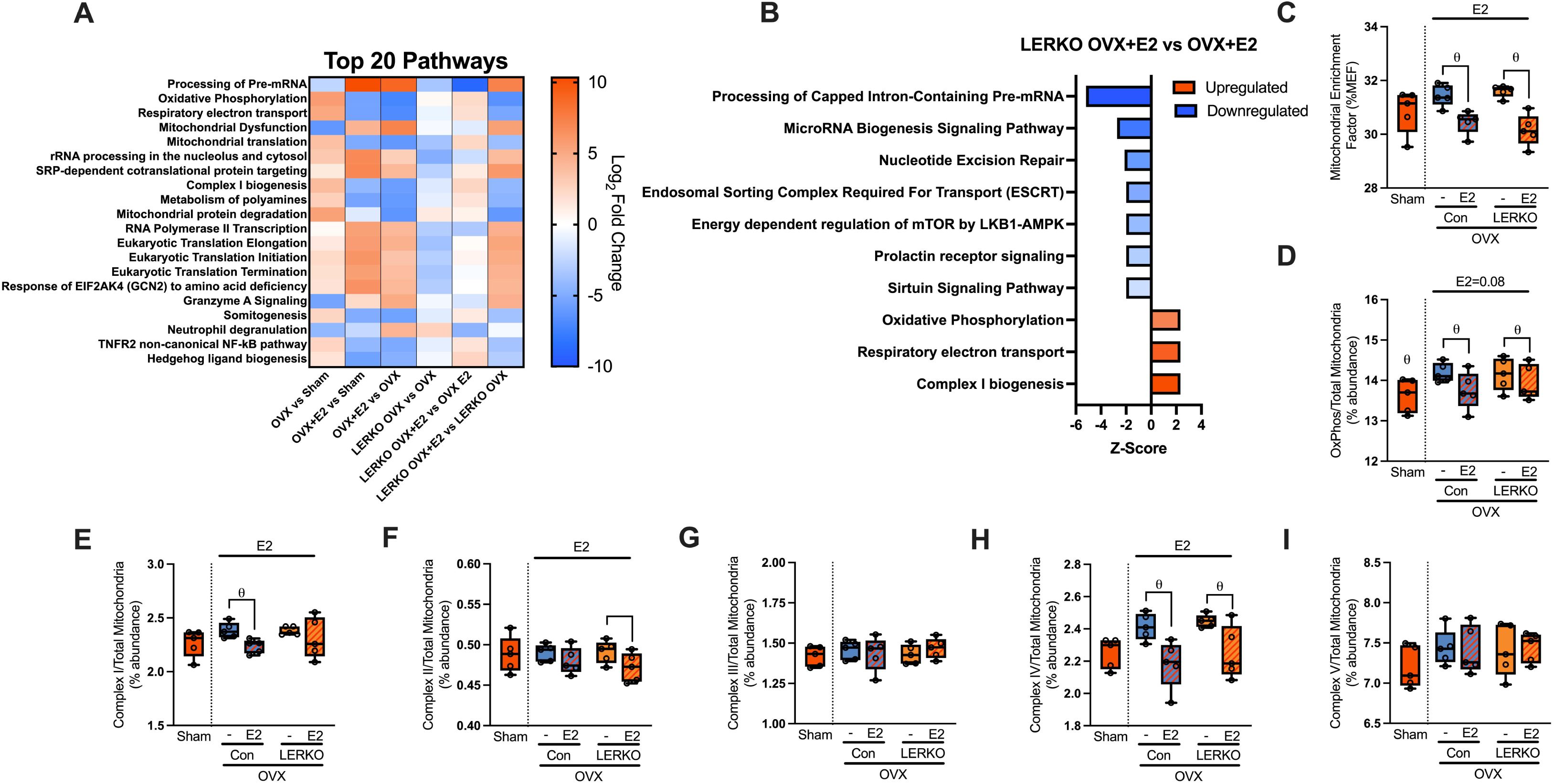
**The *hepatic mitochondrial proteome is dynamically altered by E2 replacement.*** Comparison analysis showing the top 20 altered pathways across all assessed groups (A). Top 10 significant pathways between LERKO OVX+E2 versus Con OVX+E2 mice (B). Mitochondrial enrichment factor (%MEF) was calculated by dividing the total mitochondrial protein expression by the summed expression of the entire proteome for each sample (C). Expression of OxPhos proteins normalized to the mitochondrial proteome (D). Individual Complex I (E), II (F), III (G), IV (H), V (I) expression within the mitochondrial proteome. All pathway analysis was performed with significant set at Z-score > ± 2 and Log_10_ p-value > 1.3. Data are represented as means ± SE (n=5 per group). E2 represents main effect of E2 replacement following Two-way ANOVA. Θ depicts significant unpaired t-test between Sham and Con OVX mice. Significance was set to p < 0.05.

Given our previous findings that OVX robustly impacts hepatic mitochondrial function,^38^ we examined mitochondrial content and proteins involved in energy transduction. Referencing MitoCarta3.0, we determined the mitochondrial enrichment factor (MEF) across all samples by correcting mitochondrial protein MS2 intensity against the total proteome. This revealed an E2-mediated reduction in the overall expression of mitochondrial proteins in both Con and LERKO OVX mice (Fig. 2C, p<0.0001). However, E2 replacement tended to lower OxPhos expression to comparable levels observed in Sham mice (Fig. 2D, p=0.08). Further assessment of individual complex protein expression revealed consistent E2-mediated reductions in the expression of Complex I (Fig. 2E, p<0.05), II (Fig. 2F, p<0.05), and IV (Fig. 2H, p<0.01) in both Con and LERKO mice. Complexes III and V were unaltered by E2 replacement or hepatic ERα expression. Importantly, hepatic ERα expression did not influence either total OxPhos or individual complex protein expression, suggesting the purported increase in OxPhos in LERKO OVX+E2 identified through pathway analysis is likely attributed to heightened levels of OxPhos assembly factors.

### ERα-mediated modulation of hepatic mitochondrial respiratory capacity

While we have previously reported reduced mitochondrial respiratory coupling control in OVX mice^18^, it remains unknown whether E2 replacement following OVX directly regulates hepatic mitochondrial respiratory capacity. To address this, as well as whether this process is dependent on ERα-signaling, we assessed mitochondrial *J*O_2_ under palmitate-supported respiratory conditions in isolated hepatic mitochondrial fractions from Con and LERKO mice. In Con OVX mice, E2 replacement elicited enhanced mitochondrial *J*O_2_ across basal and State 3 (ADP-stimulated) respiratory states (Fig. 3A, p<0.05), despite possessing lower OxPHOS protein expression (Fig. 2D). This enhancement did not occur under the State 3 plus succinate condition. Intriguingly, E2-mediated increases in mitochondrial *J*O_2_ were absent in LERKO OVX mice, with LERKO OVX+E2 mice maintaining reduced mitochondrial *J*O_2_ across all respiratory states compared to Con OVX+E2 mice (Fig. 3A, Int: p<0.05). Notably, E2 replacement specifically reduced maximal mitochondrial respiratory capacity following the addition of FCCP in LERKO mice (Fig. 3A, p < 0.05), indicating impairments in ETS capacity.

**Figure 3.**
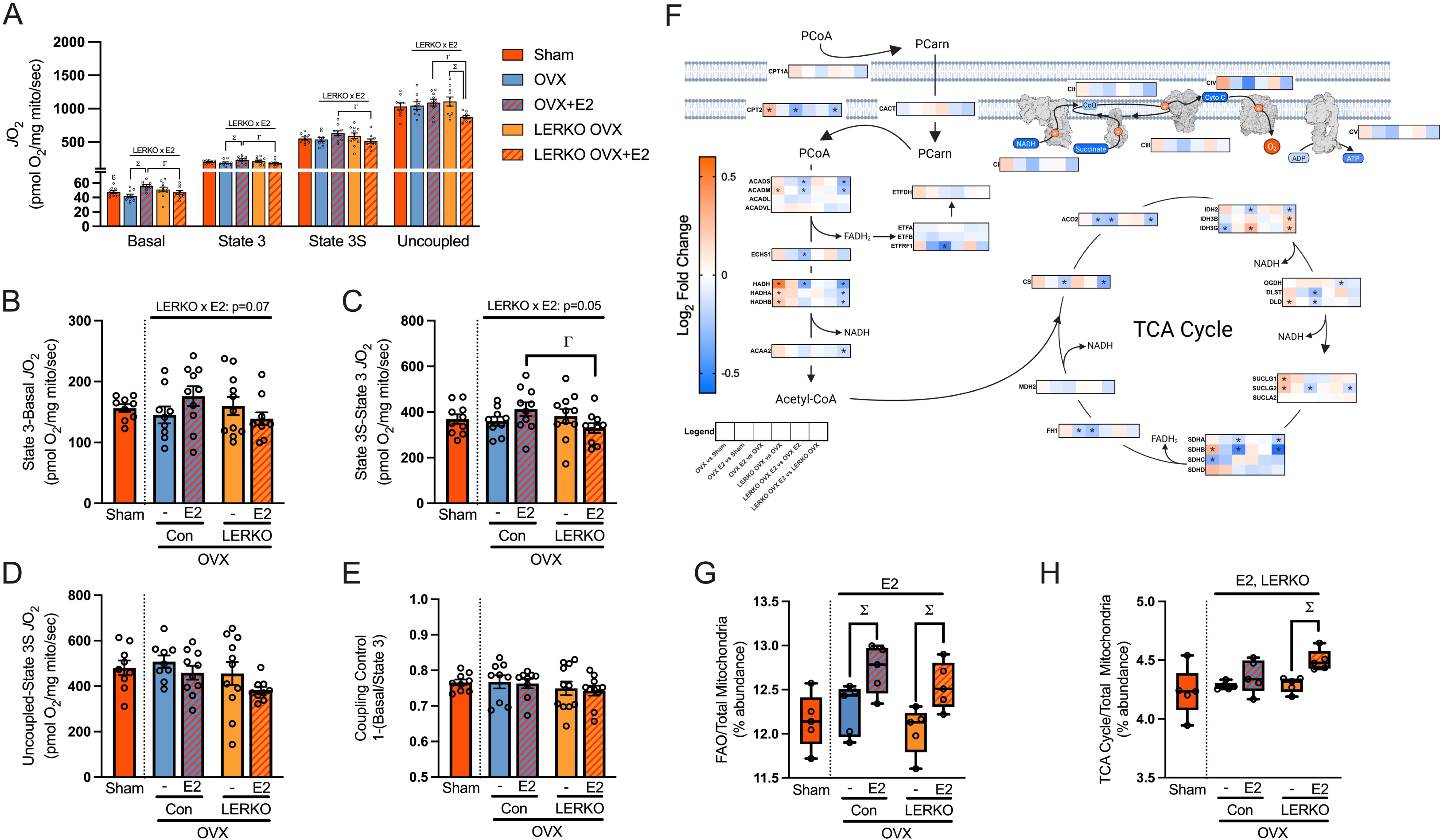
***Mitochondrial respiratory capacity is increased by E2 through hepatic ERα-signaling.*** Palmitoyl-CoA (PCoA)-supported mitochondrial oxygen consumption (*J*O_2_) across multiple respiratory states in isolated hepatic mitochondria (A). Mitochondrial response to individual substrate conditions was determined for ADP-dependent (B), succinate-dependent (C), and spare capacity (D). Mitochondrial coupling control was extrapolated from basal and State 3 *J*O_2_ (E). Proteomic log_2_ fold change in expression of fatty acid oxidation (FAO) enzymes within the mitochondria (F). Percent (%) abundance of FAO (G) and TCA cycle enzymes (H) represented within the mitochondrial proteome. Data are represented as means ± SE (n=9-11 per group in A-E; n=5 per group in G-H). LERKO x E2 signifies significant interaction between LERKO and E2 replacement following Two-way ANOVA, with Γ and Σ represent significant LSD-post hoc for LERKO and E2 replacement, respectively. ε represents significant difference between Sham and OVX+E2 replacement mice via unpaired t-test. Significance established *a priori* p < 0.05.

Importantly, differences in mitochondrial *J*O_2_ between Con OVX+E2 and LERKO OVX+E2 mice cannot fully be ascribed to differences in basal rates of mitochondrial *J*O_2._ This is shown by the Con OVX+E2 mice displaying expanded ADP-dependent (Fig. 3B, Int: p=0.07) and succinate-dependent (Fig. 3C, Int: p=0.05) mitochondrial *J*O_2_ compared to the LERKO OVX+E2 mice. No differences in spare capacity (Fig. 3D, Int: p=0.05) or mitochondrial coupling control (Fig. 3D, Int: p=0.05) were found across groups. In conjunction with coupling control ratios, no effect of LERKO or E2 replacement were apparent in mitochondrial conductance during the CK clamp (**Fig. S1**). This retention of mitochondrial coupling control further supports that deficiencies in mitochondrial respiratory capacity occur upstream of ATP synthesis.

To determine if reductions in mitochondrial *J*O_2_ in LERKO OVX+E2 mice may be attributed to lower expression of catabolic enzymes involved in fatty acid oxidation (FAO) and/or the TCA cycle, we compared the log_2_FC of all identified proteins directly involved in these processes. Mirroring total mitochondrial protein expression, these enzymes were largely lower in expression in both Con and LERKO mice receiving E2 replacement (Fig. 3F). Despite this, FAO and TCA cycle enzymes composed a greater percentage of the total mitochondrial proteome of both Con and LERKO mice receiving E2 replacement (Fig. 3G**-H**, p<0.05), with several individual proteins displaying heightened expression following E2 replacement (**Fig. S2**).

Collectively, these data are suggestive of heightening oxidative capacity through FAO and the TCA cycle in E2 replacement mice, while limitations in mitochondrial respiratory capacity apparent in LERKO OVX+E2 mice are likely attributed to reduced capacity within the ETS.

### Estrogen reduces hepatic mitochondrial H_2_O_2_ production

Alongside measures of PCoA-supported mitochondrial *J*O_2_, real-time measures of mitochondrial H_2_O_2_ emission were performed. Under basal respiratory conditions, E2 replacement promoted markedly lower H_2_O_2_ production in both Con and LERKO OVX mice (Fig. 4A, p<0.05), which was further maintained under State 3 and State 3S conditions (Fig. 4A, p<0.05). To determine % electron leak, H_2_O_2_ production was normalized to mitochondrial *J*O_2_, revealing E2 replacement diminished mitochondrial electron leak under basal (Fig. 4B, main effect p<0.05) but not State 3 or State 3S respiratory conditions (Fig. 4C**-D**). To determine if these modifications to basal mitochondrial H_2_O_2_ emission were attributed to differences in antioxidant capacity, we selectively inhibited the peroxiredoxin and glutathione antioxidant pathways to assess the raw production rates of mitochondria H_2_O_2_. Interestingly, we observed no alterations in the antioxidant capacity within either antioxidant pathway caused by E2 replacement or hepatic ERα expression (Fig. 4E**-F**). Analysis of antioxidant protein expression revealed minimal differences in mitochondrial antioxidant enzyme expression (Fig. 4G), even following correction for total mitochondrial protein expression (**Fig. S3**).

**Figure 4.**
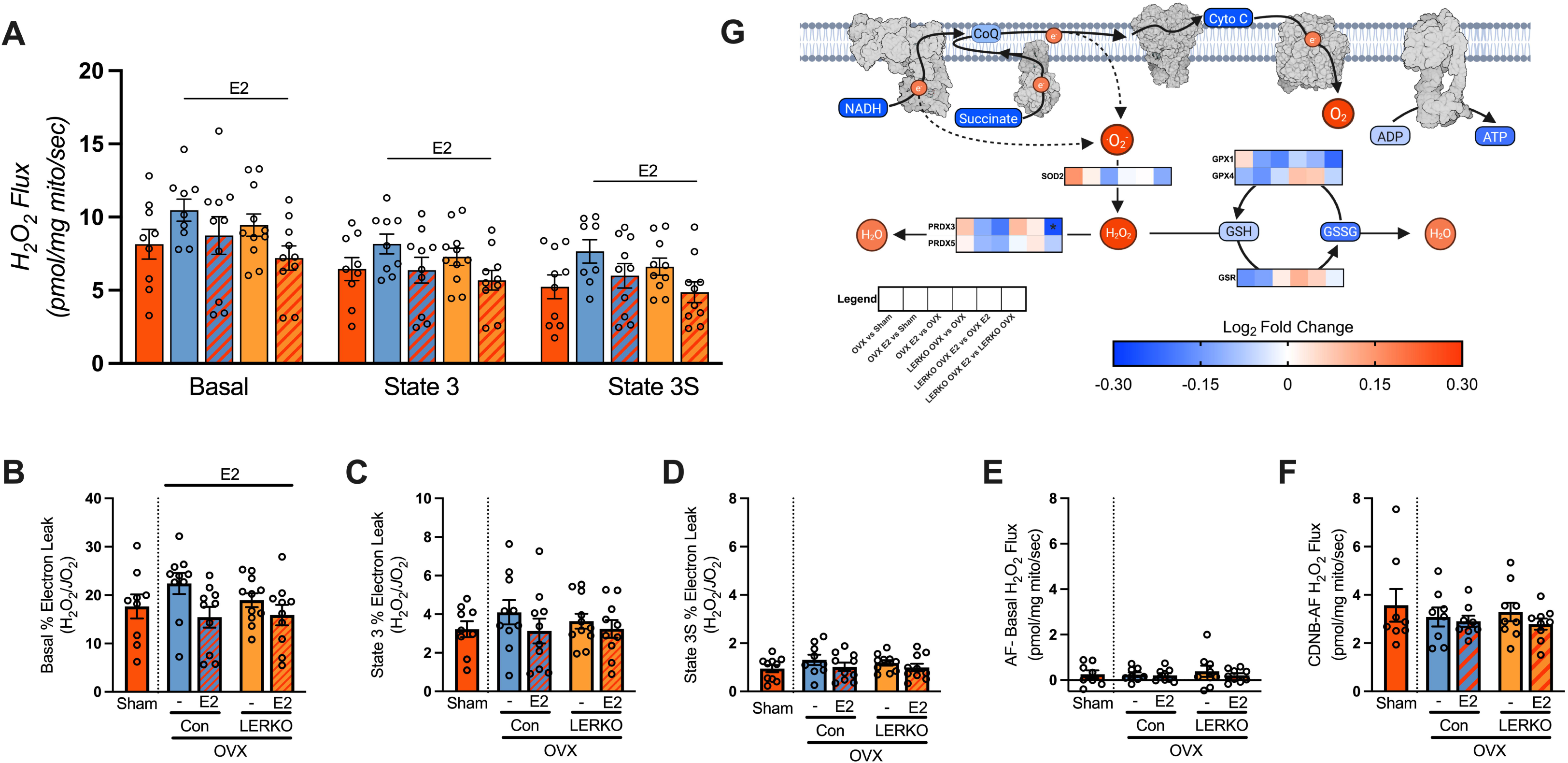
***Estrogen quenches mitochondrial-derived H_2_O_2_ production.*** Isolated mitochondrial H_2_O_2_ emission under basal, and State 3, and State 3S PCoA-supported respiratory conditions (A). Percent (%) electron leak following correction for mitochondrial oxygen consumption (*J*O_2_) under basal (B) and state 3 conditions (C). Antioxidant capacity within the peroxiredoxin (D) and glutathione antioxidant capacity (E) and total antioxidant capacity (F). Protein expression between groups was compared for major enzymes within the peroxiredoxin and glutathione antioxidant pathways (G). Data are represented as means ± SE (n=8-11 per group; A-F). Data are represented as log_2_ fold change between groups (n=5 per groups; G). E2 represents main effect of E2 replacement following Two-way ANOVA between Con OVX and LERKO OVX. Significance established *a priori* p < 0.05.

### Estrogen-mediated regulation of branched-chain keto acid catabolism

Emerging evidence suggests that diets enriched for essential amino acids, particularly BCAAs, prevent whole-body weight gain and hepatic lipid deposition through ERα-dependent signaling post-OVX^39^. To assess whether BCAA catabolism was altered by ERα signaling, we performed novel assessments of BCAA-supported mitochondrial respiration. Each BCAA is oxidized through unique enzymatic pathways, resulting in the production of acetyl-CoA (KIC), succinyl-CoA (KIV) or both metabolites (KMV). Thus, each BCAA substrate was added in series to generate an additive effect on mitochondrial respiratory capacity. Under KIC-supported (acetyl-CoA producing) basal respiration, E2 replacement promoted expansion of mitochondrial *J*O_2_ compared to Con OVX (Fig. 5A, Int: p<0.05). However, E2 replacement inversely affected LERKO OVX, resulting in decreased basal mitochondrial *J*O_2_ in LERKO OVX+E2 mice compared to both LERKO OVX and Con OVX+E2 mice (Fig. 5A, Int: p<0.01). Despite these differences in basal respiration, State 3 mitochondrial *J*O_2_ was equivalent across all groups (Fig. 5A), coinciding with no differences in ADP-dependent respiration (Fig. 5B). This resulted in Con OVX+E2 mice possessing lower mitochondrial coupling control under KIC-supported conditions (Fig. 5C, Int: p=0.05). Additionally, no differences in mitochondrial *J*O_2_ were observed following the addition of KIV (succinyl-CoA producing) (Fig. 5A). However, E2 replacement resulted in lower mitochondrial *J*O_2_ following the addition of KMV (succinyl-CoA and acetyl-CoA producing) (Fig. 5A, p<0.05) in both Con and LERKO. Similar to FAO enzymes, E2 replacement largely reduced the expression of BCAA enzymes involved in the catabolism of all 3 BCAAs (Fig. 5D). However, correction for mitochondrial protein expression revealed that BCAA enzymes composed a greater percentage of the mitochondrial proteome following E2 replacement in both Con and LERKO mice (Fig. 5E, p**<0.05)**. Interestingly, mitochondrial BCAA expression was reduced in LERKO versus Con mice. The discrepancy between BCAA-supported respiration and enzymatic expression may be due to the requirement of ATP for complete BCAA catabolism^40^, which could explain the shift from heightened to reduced respiratory rates following the addition of ADP in the E2 replacement groups.

**Figure 5.**
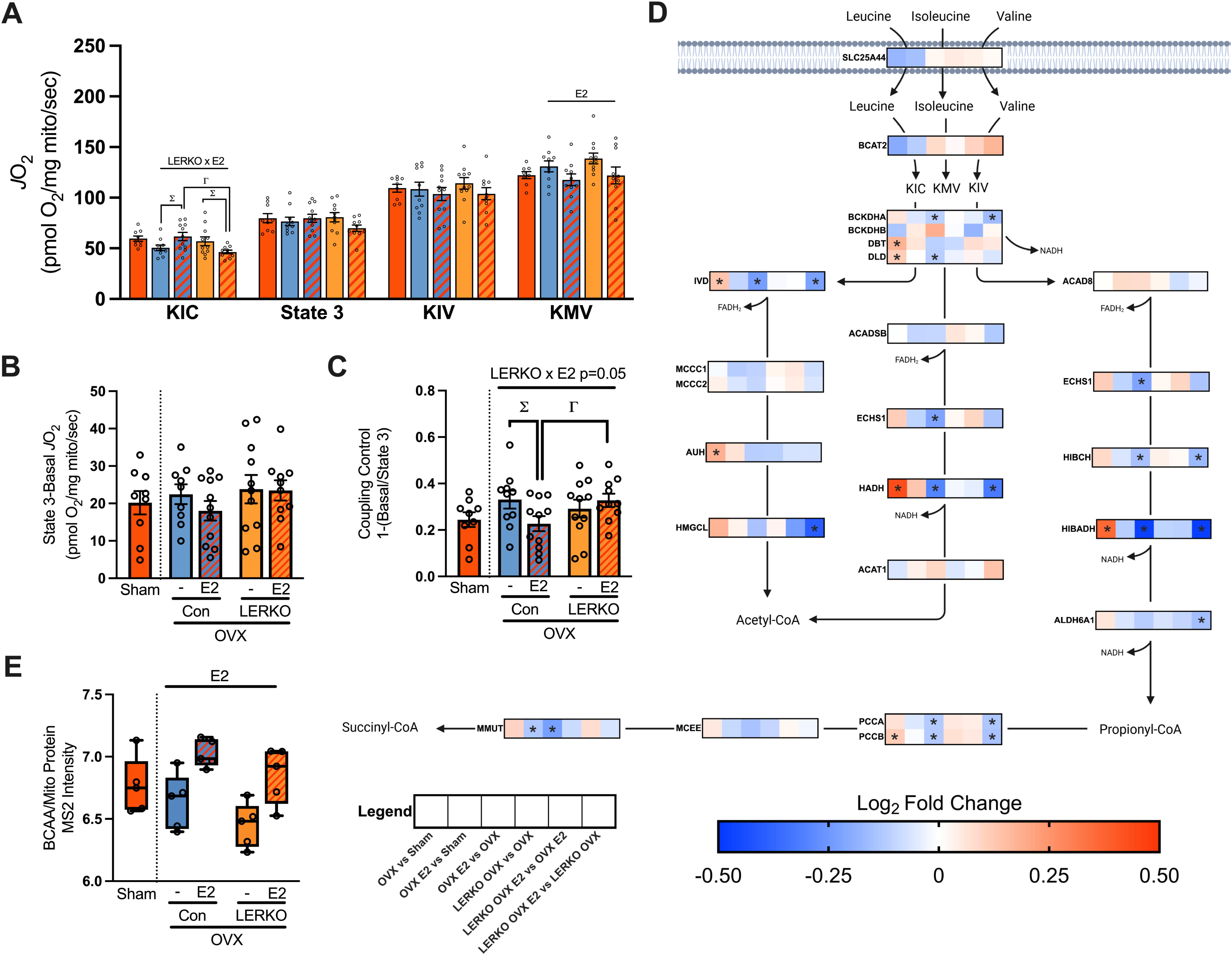
***Branched-chain keto acid (BCKA) supported mitochondrial respiration is regulated by estrogen.*** BCKA-supported mitochondrial oxygen consumption (*J*O_2_) across multiple respiratory conditions in isolated hepatic mitochondria (A). ADP-dependent respiration was determined following the addition of ADP (B). Mitochondrial coupling control was determined by the inverse relationship between basal and State 3 respiration (C). Expressional differences between groups in enzymes involved in BCAA oxidation (D), and total BCAA expression normalized to mitochondrial protein expression (E). Data are represented as means ± SE (n=9-11 per group; A-C, n=5; E). E2 represents main effect of E2 replacement using a Two-way ANOVA. LERKO x E2 signifies significant interaction between LERKO and E2 replacement following Two-way ANOVA, with Γ and Σ represent significant LSD-post hoc for LERKO and E2 replacement, respectively. Significance values are p < 0.05.

## Discussion

Although the role of ERα in modulating protection against the development of hepatic steatosis is widely debated^9,13,23,24,41^, several groups have reported that hepatic ERα is necessary for E2 replacement-mediated resistance to intrahepatic lipid deposition following a loss of ovarian function via OVX^21,22^. Previously, we have extensively phenotyped the loss of hepatic ERα in ovarian intact mice^9^. Here, we sought to determine the mitochondrial mechanisms through which hepatic ERα may modulate the therapeutic effects of E2 replacement in the treatment of hepatic steatosis following OVX. We report that E2 replacement drastically alters the hepatic proteome, resulting in a prominent reduction in mitochondrial protein expression. Despite this reduced mitochondrial content, E2 replacement promoted elevated rates of mitochondrial respiration, particularly under conditions supported by palmitate. These E2-mediated alterations in hepatic mitochondrial respiratory capacity were dependent upon hepatocyte ERα expression, with E2 replacement failing to stimulate mitochondrial respiration in LERKO mice. In contrast, E2 replacement effectively reduced hepatic mitochondrial H_2_O_2_ emission regardless of ERα expression. While we did note exacerbation of intrahepatic triglyceride levels in LERKO mice with OVX, E2-mediated reductions in hepatic steatosis were observed independent of hepatic ERα expression. Overall, the data suggest that while E2 fosters protection against hepatic steatosis irrespective of hepatic ERα, E2-mediated enhancements in mitochondrial respiratory capacity are directly modulated by hepatic ERα signaling mechanisms.

Studies utilizing chromatin immunoprecipitation (ChIP) methodologies have identified extensive and dynamic regulation of genes involved in both lipid synthesis and fatty acid oxidation (FAO) by hepatic ERα signaling^8,10,12^. Models of hepatic ERα deletion result in enriched mRNA levels of lipid biosynthesis-associated genes, indicating a prominent role of hepatic ERα in regulating lipid handling within the liver^9^. Previous investigations have suggested the protective effects of E2 against the development/treatment of hepatic steatosis function through hepatic ERα signaling^13^, particularly following OVX^21,22^. As noted, LERKO mice did have increased hepatic triglyceride content compared to control mice in all conditions and continued to be responsive to E2-mediated reductions in hepatic triglycerides, but levels still remained higher than Con OVX+E2 mice. Proteomics results confirmed E2-mediated increase in the % composition of FAO proteins within the mitochondria of both Con and LERKO livers, suggesting E2 heightened fat oxidative capacity. Three main factors likely account for the contrasting results within the present and prior reports in combined LERKO and OVX models: method of gene knockout and the timing of both OVX surgery and E2 replacement. Previous reports have relied upon the use of traditional transgenic LERKO models, in which hepatic ERα is deleted during embryonic development. Furthermore, in these studies, OVX surgery was performed in relatively young mice (<8 weeks old), with E2 replacement immediately following OVX. This combined use of transgenic models and the young age at which OVX was performed may prevent engagement of E2-mediated signaling pathways within the liver. Because of this, we utilized AAV-Cre to generate inducible hepatocyte-specific ERα knockout following OVX in middle aged mice (36-40 weeks old), permitting prolonged engagement of estrogen-mediated signaling within the liver prior to both OVX and generation of LERKO mice. Additionally, E2 replacement occurred 6-weeks post-OVX to focus our analysis on whether hepatic ERα is necessary for E2-mediated treatment of hepatic steatosis rather than a design focused on prevention in which E2 is given immediately following OVX.

Within skeletal muscle, OVX rapidly imposes deleterious effects on mitochondrial respiratory capacity. Torres et al. previously reported reductions in mitochondrial respiration 2 weeks following OVX, coinciding with a 2-fold increase in mitochondrial H_2_O_2_ production^42^. Both respiratory deficits and H_2_O_2_ production were restored with a 2-week E2 replacement^42^.

Relatively few studies have focused on hepatic mitochondrial respiration following OVX. We recently reported subtle changes to hepatic mitochondrial respiration 5 weeks post-OVX in young (16-19 week old) mice^18^. Conversely, mice in this study displayed suppressed PCoA-supported mitochondrial *J*O_2_ following OVX. Discrepancies between these studies would suggest OVX-induced mitochondrial deficiencies may take longer to manifest within the liver than skeletal muscle. Notably, no differences in either mitochondrial coupling control or conductance were identified in OVX mice, meaning alterations in mitochondrial respiration are not due to differences in the relative respiratory efficiency within the oxidative phosphorylation system. Following the addition of succinate, a complex II substrate, impairments in mitochondrial respiratory capacity were no longer apparent between Con OVX and Con OVX+E2, suggesting that E2 exerts its effects primarily through complex I. E2 replacement has been shown to increase Complex I and III activity in both liver and skeletal muscle^42,43^, coinciding with increase NADH oxidation rates in hepatic mitochondria^43^. Enhanced Complex I activity may be attributed to increased levels of flavin mononucleotide (FMN) following E2 replacement^44^. In primary hepatocytes, E2 has recently been shown to directly interact with FMN cofactor within the NADH binding site of Complex I^45^.

We have previously reported no functional impairments within hepatic mitochondria following the deletion of hepatic ERα in mice with intact ovaries^9^. In the present study, E2 replacement promoted enhanced mitochondrial respiratory capacity in Con OVX but not LERKO OVX mice indicative of a direct role for ERα-mediated signaling in modulating the beneficial effects of E2 replacement for this outcome. Similar to these results, Alves et al. has recently reported that hepatic overexpression of ERα promotes expanded hepatocyte oxygen consumption in male mice^41^. Evidence for regulation of mitochondrial respiration by ERα is not solely limited to liver, with selective activation of ERα in cardiac muscle shown to promote expanded State 3 mitochondrial *J*O_2_ following OVX^46^. ERα is known to exert transcriptional control over various mitochondrial components by binding to estrogen response elements (ERE) within the nucleus^8^. Likewise, E2 has been shown to stimulate the import of ERα (and Erβ) into the mitochondria, inducing the transcription of mitochondrial DNA (mtDNA) encoded genes, including integral components of Complexes I, III, IV, and V^47^. Together, these data suggest E2 replacement-mediated improvements in mitochondrial respiration are likely induced by ERα-dependent elevation in the transcription of mitochondrial transcription factor A (TFAM), ETS complex proteins, and mtDNA copy number. This is supported by evidence of such transcriptional alterations noted in liver^44^, primary hepatocytes^48^, and HepG2 cells following E2 treatment^49,50^. Despite this, our proteomics evidence suggests E2-mediates an unexpected lowering of mitochondrial proteins, including OxPhos proteins, in both Con and LERKO mice. We have recently identified estrogen as a potent regulator of hepatic mitophagy^34^, or the selective degradation of dysfunctional mitochondria. Thus, reductions in mitochondrial protein expression may be due to E2 activation of the mitophagy pathway, which permits the maintenance of optimally functioning pools of mitochondria. In contrast, reduced mitophagy in OVX mice allows the accrual of functionally inept mitochondria. Additionally, E2 promoted increased mitochondrial levels of FAO proteins, suggestive of enhanced capacity to catabolize fatty acid substrates and alleviate steatosis levels. Collectively, this evidence suggests that the role of ERα in modulating hepatic mitochondrial respiratory capacity may extend beyond transcriptional regulation. Further investigation is needed to fully understand the mechanisms by which E2/ERα modulates mitochondrial respiratory capacity, particularly through Complex I.

Acute E2 exposure is known to stimulate rapid mitochondrial uptake of E2, producing reductions in mitochondrial H_2_O_2_ production without alteration to mitochondrial *J*O_2_^42,51^. Here, E2 replacement effectively quenched H_2_O_2_ emission under basal and State 3 respiratory conditions, independent of hepatic ERα expression. While depletion of hepatic mitochondrial GSH and Gpx1 have been reported following OVX^52,53^, antioxidant capacity was unaltered across all groups in the present study, suggesting that E2-mediated reductions in H_2_O_2_ emissions are not attributed to changes in mitochondrial antioxidant enzymatic processes.

Likewise, unaltered liver antioxidant redox status has been reported following both OVX and OVX+E2 replacement^43^. While the antioxidant properties of E2 likely contribute to the quenching of H_2_O_2_ production in the present study, E2 has been shown to improve mitochondrial membrane integrity and increase mitochondrial membrane potential^54^. In contrast to cholesterol, membrane microviscosity is decreased by E2 within the mitochondrial inner membrane by promoting reduced lipid packing, which contributes to a more efficient transfer of electrons through ETS subunits^42^. Together, this suggests that E2 replacement may reduce mitochondrial H_2_O_2_ emission through its own antioxidant capacity and/or by improving mitochondrial membrane fluid dynamics, limiting electron leak and sustaining mitochondrial membrane potential.

In individuals with MASLD, elevated levels of both circulating and intrahepatic amino acids are commonly reported^55,56^. While women typically possess lower levels of circulating BCAAs compared to men, elevations in plasma BCAA correlate with severity of hepatic steatosis/MASLD in women^57^. In response to a short-term fast, female mice catabolize amino acids within the liver to a far greater extent than male mice through robust transcription of genes involved in amino acid transport and catabolism^11^. However, the liver’s capacity to utilize amino acids is severely blunted in the absence of estrogen, as seen in female aromatase knockout mice, which cannot synthesize estrogen^11^. Similarly, reductions in the transcription of genes involved in amino acid catabolism have recently been reported in LERKO mice^13^, suggesting amino acid catabolism is directly regulated by ERα. Indeed, our proteomics data confirm that E2 mediates the modulation of BCAA enzyme expression, and that LERKO mice, regardless of E2 treatment, possess reduced enzyme levels. Utilizing BCKAs, the downstream metabolites of BCAAs, we directly assessed BCKA-supported mitochondrial respiration. Under KIC-supported basal conditions, E2 significantly increased mitochondrial *J*O_2_ consumption only in the presence of hepatic ERα. However, due to the similarities between PCoA- and KIC-supported respiration, we cannot rule out that the observed elevation in basal respiration is not simply attributed to E2-mediated increases in complex I activity via hepatic ERα. During maximal BCKA-mediated respiratory conditions (State 3 KIC+KIV+KMV), E2 replacement universally lowered mitochondrial *J*O_2_. Following correcting for differences in basal respiration, E2-mediated reductions in maximal BCKA respiration appear to function through suppression of mitochondrial *J*O_2_. While a unique interaction between BCAA catabolism and hepatic ERα expression has been previously described^10,11,39,58^, we provide the first direct evidence that BCKA catabolism is dynamically modulated by ERα-dependent estrogen signaling within the liver. However, further investigation is needed to understand how ERα-mediated regulation of hepatic BCAA catabolism may alter susceptibility to and/or progression of hepatic steatosis in post-menopausal conditions.

There are several limitations within this study that must be noted. First, the use of OVX to induce a menopause-like state in mice results in rapid alterations in the circulating levels of various sex hormones, which does not accurately replicate the human condition. Additionally, while the use of whole-liver proteomics to show increased mitochondrial content, paired with measures of mitochondrial bioenergetics support our overall conclusion that impairments in mitochondrial respiration are due to deficits in mitochondrial respiratory capacity, the use of isolated mitochondrial isolates for proteomics would have provided a more direct explanation for the observed respiratory deficits in Con OVX and LERKO OVX+E2 mice. Although we sought to understand the role of estrogen in modulating the susceptibility to hepatic steatosis, the inclusion of low-fat diet fed groups would have allowed for greater understanding of the direct effects of E2 on hepatic mitochondrial bioenergetics, independent of the compounding effects of HFD feeding.

This study builds upon previous work suggesting that hepatic ERα is necessary for E2-mediated protection against hepatic steatosis following OVX^21,22^. Here, we report that E2 restores protection against hepatic steatosis irrespective of ERα expression, coinciding with E2-mediated increased in mitochondrial FAO enzymes. Existing literature would suggest that E2’s capacity to positively influence metabolism in adipose and muscle also likely contributed to the lowering of hepatic triglycerides. However, E2-mediated improvements to mitochondrial respiratory capacity occur in an ERα-dependent manner following OVX, suggesting a direct role of Erα signaling in modulating hepatic mitochondrial respiration. At the same time, E2 treatment lowered mitochondrial H_2_O_2_ emission independent of hepatic ERα expression. Finally, we identify E2-mediated regulation of hepatic BCKA-supported mitochondrial respiration, suggesting ERα may exert control over BCKA catabolism within the liver. Based on the data obtained by the present study, targeting the selective activation of hepatic ERα may serve as a novel therapeutic target for improving hepatic mitochondrial bioenergetics. Future studies should aim to further interrogate the mechanistic regulation of hepatic mitochondrial bioenergetics by E2 and hepatic ERα in the context of MASLD treatment in post-menopausal women.

## Acknowledgements and Funding

This study was supported by VA Merit Review grant 1I01BX002567 (JPT), NIH P20GM144269 (CSM, EMM, JPT), NIH T32DK128770 (EF, SFS) and NIH T32AG07811 (BAK).

## Author Contributions

Acquisition of funding (JPT), Investigation (EF, BAK, SFS, JAA, CSM, EMM), Resources and Methodology (EF, BAK), Supervision (JPT), Writing – Original Draft Preparation (EF, JPT), Writing – Review & Editing (all authors)

## Statements and Declarations

The authors declare no competing interest.

---

**Figure S1.** Graphical depiction of chemical concentrations during creatine kinase (CK) clamp (A). Mitochondrial *J*O_2_ during the CK clamp (B). Mitochondrial conductance calculated from the linear relationship between mitochondrial *J*O_2_ and ΔG_ATP_ (C). Data are represented as means ± SE (n=9-11 per group. Significance established *a priori* p < 0.05.

**Figure S2.** Fatty acid oxidation proteins with significantly altered protein expression following normalization to mitochondrial protein expression, including SLC25A20A (A), CPT2 (B), ACADL (C), ACADVL (D), ETFA (E), ETFB (F), ETFD (G). Data are represented as means ± SE (n=5 per group). E2 and LERKO represent significant main effect of E2 replacement or LERKO, respectively, via Two-way ANOVA. Σ represents significant LSD post-hoc for E2 replacement. ε and Θ represent significant unpaired t-test between Sham and Con OVX+E2 and Sham and Con OVX, respectively. Significance established *a priori* p < 0.05.

**Figure S3.** Branched-chain amino acid catabolic enzymes with significantly altered protein expression following normalization to mitochondrial protein expression, including SLC25A44 (A), BCKDH (B), MCC (C), ACAD8 (D), HIBADH (E), ALDH6A1 (F), PCC (G) and MMUT (H). Data are represented as means ± SE (n=5 per group). E2 and LERKO represent significant main effect of E2 replacement or LERKO, respectively, via Two-way ANOVA. Σ and Γ represents significant LSD post-hoc for E2 replacement and LERKO, respectively. Θ represents significant unpaired t-test between Sham and Con OVX. Significance established *a priori* p < 0.05.

## References

1 Cui, J., Shen, Y. & Li, R. Estrogen synthesis and signaling pathways during aging: from periphery to brain. Trends Mol Med 19, 197–209 (2013). 10.1016/j.molmed.2012.12.007

2 Zhu, B. T., Han, G.-Z., Shim, J.-Y., Wen, Y. & Jiang, X.-R. Quantitative Structure-Activity Relationship of Various Endogenous Estrogen Metabolites for Human Estrogen Receptor α and β Subtypes: Insights into the Structural Determinants Favoring a Differential Subtype Binding. Endocrinology 147, 4132–4150 (2006). 10.1210/en.2006-0113

3 Blair, R. M. et al. The estrogen receptor relative binding affinities of 188 natural and xenochemicals: structural diversity of ligands. Toxicol Sci 54, 138–153 (2000). 10.1093/toxsci/54.1.138

4 Filardo, E. J., Quinn, J. A., Frackelton, A. R., Jr. & Bland, K. I. Estrogen action via the G protein-coupled receptor, GPR30: stimulation of adenylyl cyclase and cAMP-mediated attenuation of the epidermal growth factor receptor-to-MAPK signaling axis. Mol Endocrinol 16, 70–84 (2002). 10.1210/mend.16.1.0758

5 Hevener, A. L., Ribas, V., Moore, T. M. & Zhou, Z. The Impact of Skeletal Muscle ERα on Mitochondrial Function and Metabolic Health. Endocrinology 161 (2020). 10.1210/endocr/bqz017

6 Yang, X. et al. Tissue-specific expression and regulation of sexually dimorphic genes in mice. Genome research 16, 995–1004 (2006).

7 Zhang, Y. et al. Transcriptional profiling of human liver identifies sex-biased genes associated with polygenic dyslipidemia and coronary artery disease. PLoS One 6, e23506 (2011). 10.1371/journal.pone.0023506

8 Gao, H., Falt, S., Sandelin, A., Gustafsson, J. A. & Dahlman-Wright, K. Genome-wide identification of estrogen receptor alpha-binding sites in mouse liver. Mol Endocrinol 22, 10–22 (2008). 10.1210/me.2007-0121

9 Fuller, K. N. Z. et al. Pre– and Post–Sexual Maturity Liver-specific ERα Knockout Does Not Impact Hepatic Mitochondrial Function. Journal of the Endocrine Society 7 (2023). 10.1210/jendso/bvad053

10 Della Torre, S. et al. An Essential Role for Liver ERα in Coupling Hepatic Metabolism to the Reproductive Cycle. Cell Reports 15, 360–371 (2016). 10.1016/j.celrep.2016.03.019

11 Della Torre, S. et al. Short-Term Fasting Reveals Amino Acid Metabolism as a Major Sex-Discriminating Factor in the Liver. Cell Metab 28, 256–267.e255 (2018). 10.1016/j.cmet.2018.05.021

12 Villa, A. et al. Tetradian oscillation of estrogen receptor α is necessary to prevent liver lipid deposition. Proc Natl Acad Sci U S A 109, 11806–11811 (2012). 10.1073/pnas.1205797109

13 Meda, C. et al. Hepatic ERα accounts for sex differences in the ability to cope with an excess of dietary lipids. Molecular Metabolism 32, 97–108 (2020). 10.1016/j.molmet.2019.12.009

14 Lonardo, A. et al. Sex Differences in Nonalcoholic Fatty Liver Disease: State of the Art and Identification of Research Gaps. Hepatology 70, 1457–1469 (2019). 10.1002/hep.30626

15 PARK, S. H., et al. Prevalence and risk factors of non-alcoholic fatty liver disease among Korean adults. Journal of Gastroenterology and Hepatology 21, 138–143 (2006). 10.1111/j.1440-1746.2005.04086.x

16 Long, M. T. et al. A simple clinical model predicts incident hepatic steatosis in a community-based cohort: The Framingham Heart Study. Liver International 38, 1495–1503 (2018). 10.1111/liv.13709

17 Wang, Z., Xu, M., Hu, Z., Hultström, M. & Lai, E. Sex-specific prevalence of fatty liver disease and associated metabolic factors in Wuhan, south central China. European Journal of Gastroenterology & Hepatology 26, 1015–1021 (2014). 10.1097/meg.0000000000000151

18 Fuller, K. N. Z. et al. Estradiol treatment or modest exercise improves hepatic health and mitochondrial outcomes in female mice following ovariectomy. American Journal of Physiology-Endocrinology and Metabolism 320, E1020–E1031 (2021). 10.1152/ajpendo.00013.2021

19 Kumari, R. et al. VCD-induced menopause mouse model reveals reprogramming of hepatic metabolism. Mol Metab 82, 101908 (2024). 10.1016/j.molmet.2024.101908

20 Clark, J. M., Brancati, F. L. & Diehl, A. M. Nonalcoholic fatty liver disease. Gastroenterology 122, 1649–1657 (2002). 10.1053/gast.2002.33573

21 Zhu, L. et al. Estrogen Treatment After Ovariectomy Protects Against Fatty Liver and May Improve Pathway-Selective Insulin Resistance. Diabetes 62, 424–434 (2013). 10.2337/db11-1718

22 Guillaume, M. et al. Selective Liver Estrogen Receptor α Modulation Prevents Steatosis, Diabetes, and Obesity Through the Anorectic Growth Differentiation Factor 15 Hepatokine in Mice. Hepatology Communications 3, 908–924 (2019). 10.1002/hep4.1363

23 Hart-Unger, S. et al. Hormone signaling and fatty liver in females: analysis of estrogen receptor α mutant mice. International Journal of Obesity 41, 945–954 (2017). 10.1038/ijo.2017.50

24 Zhu, L. et al. Hepatocyte estrogen receptor alpha mediates estrogen action to promote reverse cholesterol transport during Western-type diet feeding. Mol Metab 8, 106–116 (2018). 10.1016/j.molmet.2017.12.012

25 Koliaki, C. et al. Adaptation of Hepatic Mitochondrial Function in Humans with Non-Alcoholic Fatty Liver Is Lost in Steatohepatitis. Cell Metabolism 21, 739–746 (2015). 10.1016/j.cmet.2015.04.004

26 Von Schulze, A. et al. Hepatic mitochondrial adaptations to physical activity: impact of sexual dimorphism, PGC1α and BNIP3-mediated mitophagy. *J Physiol* **596**, 6157-6171 (2018). 10.1113/jp276539

27 McCoin, C. S. et al. Sex modulates hepatic mitochondrial adaptations to high-fat diet and physical activity. Am J Physiol Endocrinol Metab 317, E298–E311 (2019). 10.1152/ajpendo.00098.2019

28 Fuller, K. N. Z. et al. Sex and BNIP3 genotype, rather than acute lipid injection, modulate hepatic mitochondrial function and steatosis risk in mice. J Appl Physiol (1985) 128, 1251–1261 (2020). 10.1152/japplphysiol.00035.2020

29 Fuller, K. N. Z. et al. Pre- and Post-Sexual Maturity Liver-specific ERalpha Knockout Does Not Impact Hepatic Mitochondrial Function. J Endocr Soc 7, bvad053 (2023). 10.1210/jendso/bvad053

30 Ingberg, E., Theodorsson, A., Theodorsson, E. & Strom, J. O. Methods for long-term 17β-estradiol administration to mice. Gen Comp Endocrinol 175, 188–193 (2012). 10.1016/j.ygcen.2011.11.014

31 Fuller, K. N. Z. et al. Oral combined contraceptives induce liver mitochondrial reactive oxygen species and whole-body metabolic adaptations in female mice. J Physiol 600, 5215–5245 (2022). 10.1113/jp283733

32 Morris, E. M. et al. PGC-1α overexpression results in increased hepatic fatty acid oxidation with reduced triacylglycerol accumulation and secretion. American Journal of Physiology-Gastrointestinal and Liver Physiology 303, G979–G992 (2012). 10.1152/ajpgi.00169.2012

33 Krumschnabel, G. et al. Simultaneous high-resolution measurement of mitochondrial respiration and hydrogen peroxide production. Methods Mol Biol 1264, 245–261 (2015). 10.1007/978-1-4939-2257-4_22

34 Franczak, E. et al. Loss of ovarian function prevents exercise-induced activation of hepatic mitophagic flux. Am J Physiol Endocrinol Metab 328, E869–e884 (2025). 10.1152/ajpendo.00107.2025

35 Glancy, B., Barstow, T. & Willis, W. T. Linear relation between time constant of oxygen uptake kinetics, total creatine, and mitochondrial content in vitro. American Journal of Physiology-Cell Physiology 294, C79–C87 (2008). 10.1152/ajpcell.00138.2007

36 Fisher-Wellman, K. H. et al. Mitochondrial Diagnostics: A Multiplexed Assay Platform for Comprehensive Assessment of Mitochondrial Energy Fluxes. Cell Rep 24, 3593–3606.e3510 (2018). 10.1016/j.celrep.2018.08.091

37 Rath, S. et al. MitoCarta3.0: an updated mitochondrial proteome now with sub-organelle localization and pathway annotations. Nucleic Acids Res 49, D1541–d1547 (2021). 10.1093/nar/gkaa1011

38 Fuller, K. N. Z. et al. Estradiol treatment or modest exercise improves hepatic health and mitochondrial outcomes in female mice following ovariectomy. Am J Physiol Endocrinol Metab 320, E1020–E1031 (2021). 10.1152/ajpendo.00013.2021

39 Della Torre, S. et al. Dietary essential amino acids restore liver metabolism in ovariectomized mice via hepatic estrogen receptor α. Nat Commun 12, 6883 (2021). 10.1038/s41467-021-27272-x

40 Goldberg, E. J. et al. Tissue-specific characterization of mitochondrial branched-chain keto acid oxidation using a multiplexed assay platform. Biochem J 476, 1521–1537 (2019). 10.1042/bcj20190182

41 Alves, E. S. et al. Hepatic Estrogen Receptor Alpha Overexpression Protects Against Hepatic Insulin Resistance and MASLD. Pathophysiology 32, 1 (2025).

42 Torres, M. J. et al. 17β-Estradiol Directly Lowers Mitochondrial Membrane Microviscosity and Improves Bioenergetic Function in Skeletal Muscle. Cell Metab 27, 167–179.e167 (2018). 10.1016/j.cmet.2017.10.003

43 Torres, M. J., Ryan, T. E., Lin, C. T., Zeczycki, T. N. & Neufer, P. D. Impact of 17β-estradiol on complex I kinetics and H(2)O(2) production in liver and skeletal muscle mitochondria. J Biol Chem 293, 16889–16898 (2018). 10.1074/jbc.RA118.005148

44 Tian, Y., Hong, X., Xie, Y., Guo, Z. & Yu, Q. 17β-Estradiol (E2) Upregulates the ERα/SIRT1/PGC-1α Signaling Pathway and Protects Mitochondrial Function to Prevent Bilateral Oophorectomy (OVX)-Induced Nonalcoholic Fatty Liver Disease (NAFLD). Antioxidants 12, 2100 (2023).

45 Moreira, P. I., Custódio, J., Moreno, A., Oliveira, C. R. & Santos, M. S. Tamoxifen and Estradiol Interact with the Flavin Mononucleotide Site of Complex I Leading to Mitochondrial Failure*. Journal of Biological Chemistry 281, 10143–10152 (2006). 10.1074/jbc.M510249200

46 Hamilton, D. J. et al. Estrogen receptor alpha activation enhances mitochondrial function and systemic metabolism in high-fat-fed ovariectomized mice. Physiol Rep 4 (2016). 10.14814/phy2.12913

47 Chen, J. Q., Delannoy, M., Cooke, C. & Yager, J. D. Mitochondrial localization of ERalpha and ERbeta in human MCF7 cells. Am J Physiol Endocrinol Metab 286, E1011–1022 (2004). 10.1152/ajpendo.00508.2003

48 Chen, J. et al. Enhanced Mitochondrial Gene Transcript, ATP, Bcl-2 Protein Levels, and Altered Glutathione Distribution in Ethinyl Estradiol-Treated Cultured Female Rat Hepatocytes. Toxicological Sciences 75, 271–278 (2003). 10.1093/toxsci/kfg183

49 Galmés-Pascual, B. M. et al. 17β-estradiol ameliorates lipotoxicity-induced hepatic mitochondrial oxidative stress and insulin resistance. Free Radical Biology and Medicine 150, 148–160 (2020). 10.1016/j.freeradbiomed.2020.02.016

50 Chen, J., Li, Y., Lavigne, J. A., Trush, M. A. & Yager, J. D. Increased mitochondrial superoxide production in rat liver mitochondria, rat hepatocytes, and HepG2 cells following ethinyl estradiol treatment. Toxicol Sci 51, 224–235 (1999). 10.1093/toxsci/51.2.224

51 Moats, R. K., II & Ramirez, V. D. Rapid Uptake and Binding of Estradiol-17β-6-(O-carboxymethyl)oxime:125I-labeled Bsa by Female Rat Liver1. Biology of Reproduction 58, 531–538 (1998). 10.1095/biolreprod58.2.531

52 Borrás, C. et al. Mitochondria from females exhibit higher antioxidant gene expression and lower oxidative damage than males. Free Radic Biol Med 34, 546–552 (2003). 10.1016/s0891-5849(02)01356-4

53 Valencia, A. P. et al. The presence of the ovary prevents hepatic mitochondrial oxidative stress in young and aged female mice through glutathione peroxidase 1. Experimental Gerontology 73, 14–22 (2016). 10.1016/j.exger.2015.11.011

54 Borrás, C., Gambini, J., López-Grueso, R., Pallardó, F. V. & Viña, J. Direct antioxidant and protective effect of estradiol on isolated mitochondria. Biochimica et Biophysica Acta (BBA) - Molecular Basis of Disease 1802, 205–211 (2010). 10.1016/j.bbadis.2009.09.007

55 Gaggini, M. et al. Altered amino acid concentrations in NAFLD: Impact of obesity and insulin resistance. Hepatology 67, 145–158 (2018). 10.1002/hep.29465

56 Lake, A. D. et al. Branched chain amino acid metabolism profiles in progressive human nonalcoholic fatty liver disease. Amino Acids 47, 603–615 (2015). 10.1007/s00726-014-1894-9

57 Grzych, G. et al. Plasma BCAA Changes in Patients With NAFLD Are Sex Dependent. J Clin Endocrinol Metab 105 (2020). 10.1210/clinem/dgaa175

58 Della Torre, S. et al. Amino acid-dependent activation of liver estrogen receptor alpha integrates metabolic and reproductive functions via IGF-1. Cell Metab 13, 205–214 (2011). 10.1016/j.cmet.2011.01.002

