## Supplementary figures and images for "Hepatocyte Estrogen Receptor α Mediates Estrogen-induced Augmentation of Hepatic Mitochondrial Respiration Following Ovariectomy"

### Supplemental Fig. 1

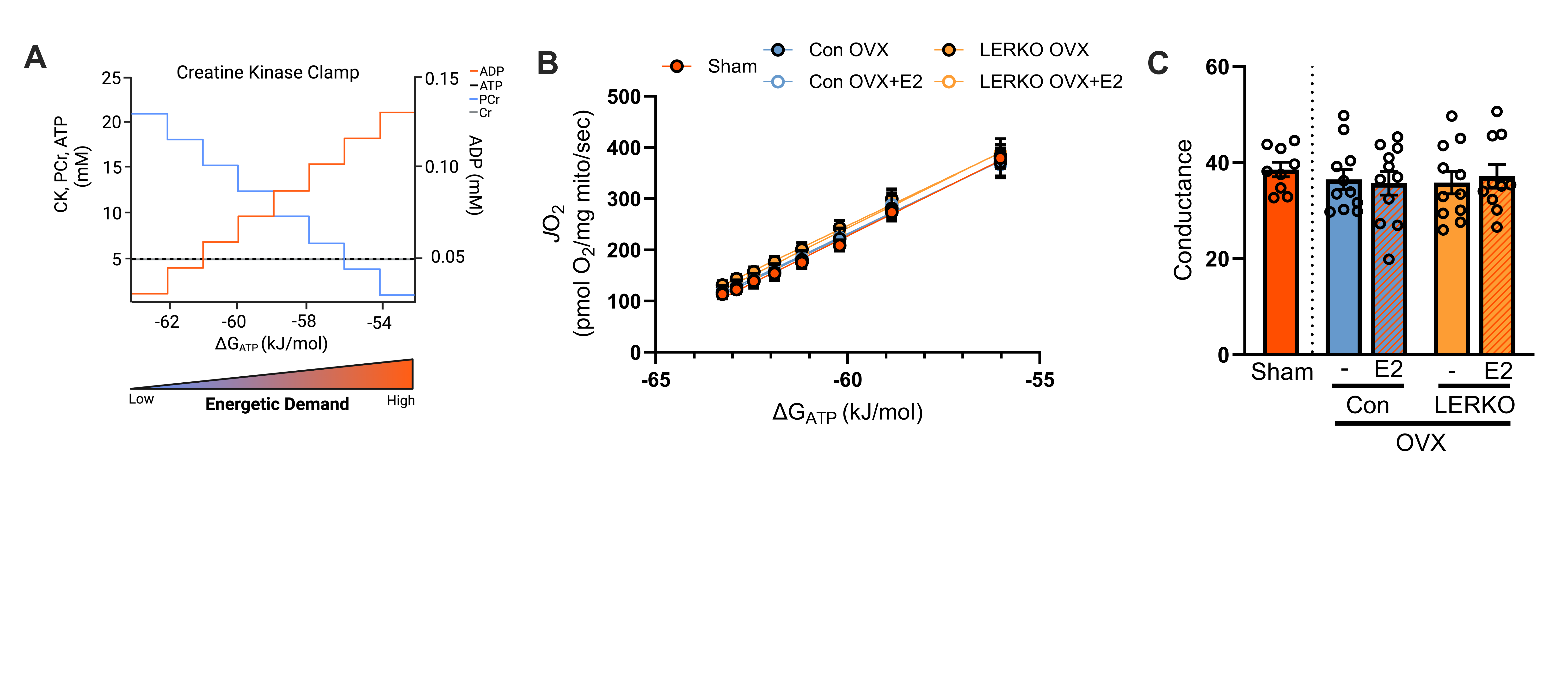

### Supplemental Fig. 2

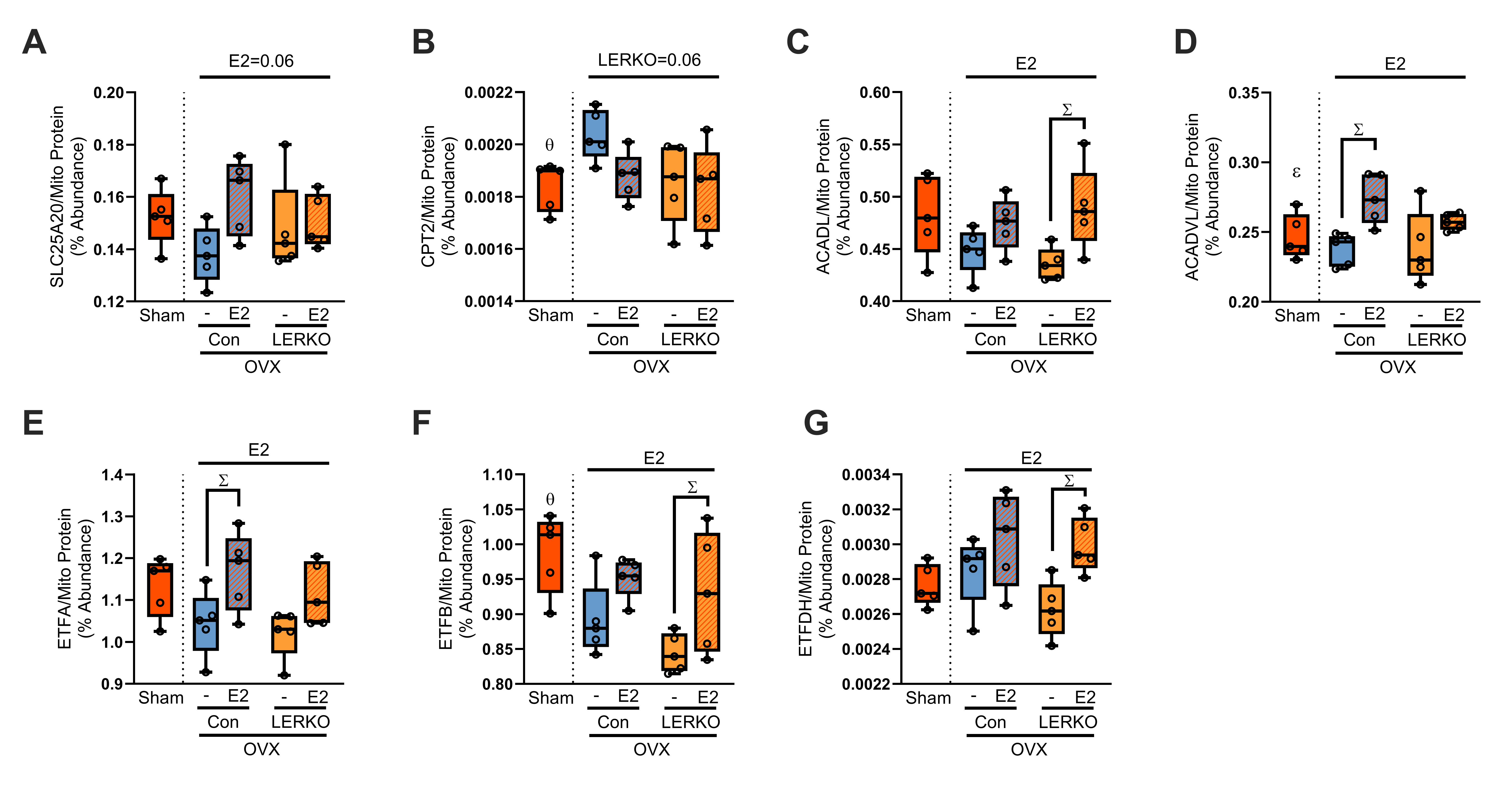

### Supplemental Fig. 3

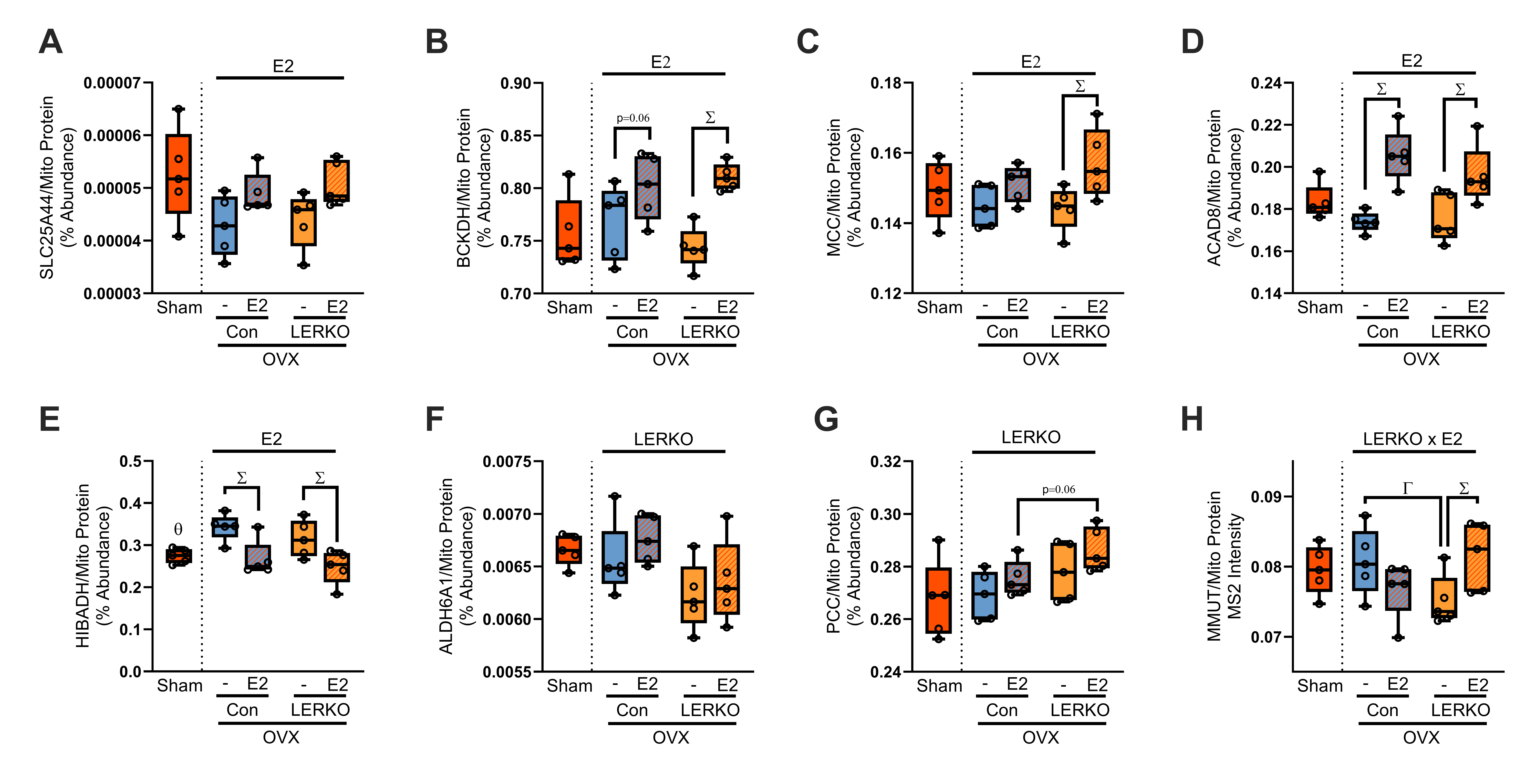
